# Repositioning of polyubiquitin alters the pathologic tau filament structure

**DOI:** 10.1101/2025.05.02.651930

**Authors:** Ryohei Watanabe, Benjamin C. Creekmore, Nabil F. Darwich, Hong Xu, Lakshmi Changolkar, Kevt’her Hoxha, Bin Zhang, Caroline M. O’Rourke, George M. Burslem, Virginia M.-Y. Lee, Yi-Wei Chang, Edward B. Lee

## Abstract

Structurally diverse tau filaments form proteinaceous aggregates in a heterogeneous group of neurodegenerative diseases called tauopathies^1^. The factors extrinsic to the highly ordered core structure that influence tau filament stability are not well understood. Here, we found that polyubiquitinated tau filaments from Alzheimer’s disease and vacuolar tauopathy human brain tissue exhibit distinct seeding patterns in mice, in association with differences in tau filament ultrastructure determined by cryo-electron microscopy. Interestingly, chemical modulation of the polarity of polyubiquitin adjacent to the tau core with the small molecule ubistatin B resulted in the repositioning of poorly structured densities towards positively charged residues on the highly structured core filament, leading to shifting of the protofilament-protofilament interface of certain vacuolar tauopathy tau filaments. These results suggest that the structure of tau filaments that are associated with different seeding activities *in vivo* can be influenced by post-translational modifications.

## Main text

Tauopathies are a heterogeneous group of aging-related neurodegenerative diseases with considerable clinical, pathological, and biochemical diversity. In these diseases, the accumulation of proteinaceous aggregates comprised of insoluble tau filaments is tightly associated with neurodegeneration. The development of cryo-electron microscopy (cryo-EM) helical reconstruction has enabled the characterization of near-atomic core structures of tau amyloid filaments from postmortem human tauopathy brains^2^ where different tau amyloid core structures are associated with different tauopathy subtypes ^3–7^. This diversity of tau filament structures is thought to result in distinct patterns of cell-to-cell propagation of tau through the brain connectome *in vivo*, representing a structural correlate to the regional and cellular patterns of selective vulnerability that define each tauopathy subtype^8,9^.

Given the heterogeneity of tau filaments across different tauopathies, elucidation of the factors that affect tau fibril stabilization may improve our understanding of mechanisms underlying tau-mediated neurodegeneration. In particular, cryo-EM density maps have demonstrated that there are additional densities adjacent to the core of tau filaments, densities which have been difficult to characterize due to their flexibility but have been speculated to correspond to post-translational modifications (PTMs) or other cofactors^10^. The effect of these extra densities on the stability of tau filaments is not well understood.

Here, we found that ubiquitinated tau filaments extracted from Alzheimer’s disease (AD) or vacuolar tauopathy (VT) brain tissues exhibit different patterns of tau seeding *in vivo* which appears to mimic the distribution and morphology of tau aggregates in each disease. This difference in biological activity was associated with differences in tau core structures including a diverse assortment of five tau filaments in VT. Using a chemical biology approach, treating tau filaments with ubistatin B (ubiB) to alter the polarity of polyubiquitin allowed for the localization of ubiquitin on tau core filaments. Surprisingly, chemical manipulation of polyubiquitin with ubiB resulted in the shifting of protofilaments in certain VT tau filaments, suggesting that post-translational modifications can affect tau fibril stability.

### Biological activity of AD and VT tau

VT is caused by mutations in VCP, a AAA+ ATPase which biochemically unfolds structured, polyubiquitinated protein substrates^11^. To demonstrate that pathological tau protein is polyubiquitinated in both AD and VT brains, sarkosyl-insoluble tau filaments were extracted from the frontal neocortices of three AD cases and three VT cases (Extended Data Tables 1 and 2). The neuropathologic description of these three VT cases has recently been reported and includes neurodegeneration of frontal and temporal lobes in association with the accumulation of tau aggregates (manuscript under review). Sarkosyl-insoluble tau was denatured with SDS, immunoprecipitated with an N-terminal tau antibody (Tau-12) ^12^, and then immunoblotted with antibodies that recognize total tau (17025) or ubiquitin (E4I2J; Fig. 1a). AD and VT tau exhibited similar banding patterns, consistent with the fact that both diseases accumulate both 3-repeat and 4-repeat tau isoforms as previously described^11^. In addition, although there was some variability, higher molecular-weight protein present on anti-ubiquitin immunoblots indicated that polyubiquitination of insoluble tau was readily detectable in both AD and VT.

**Fig. 1.**
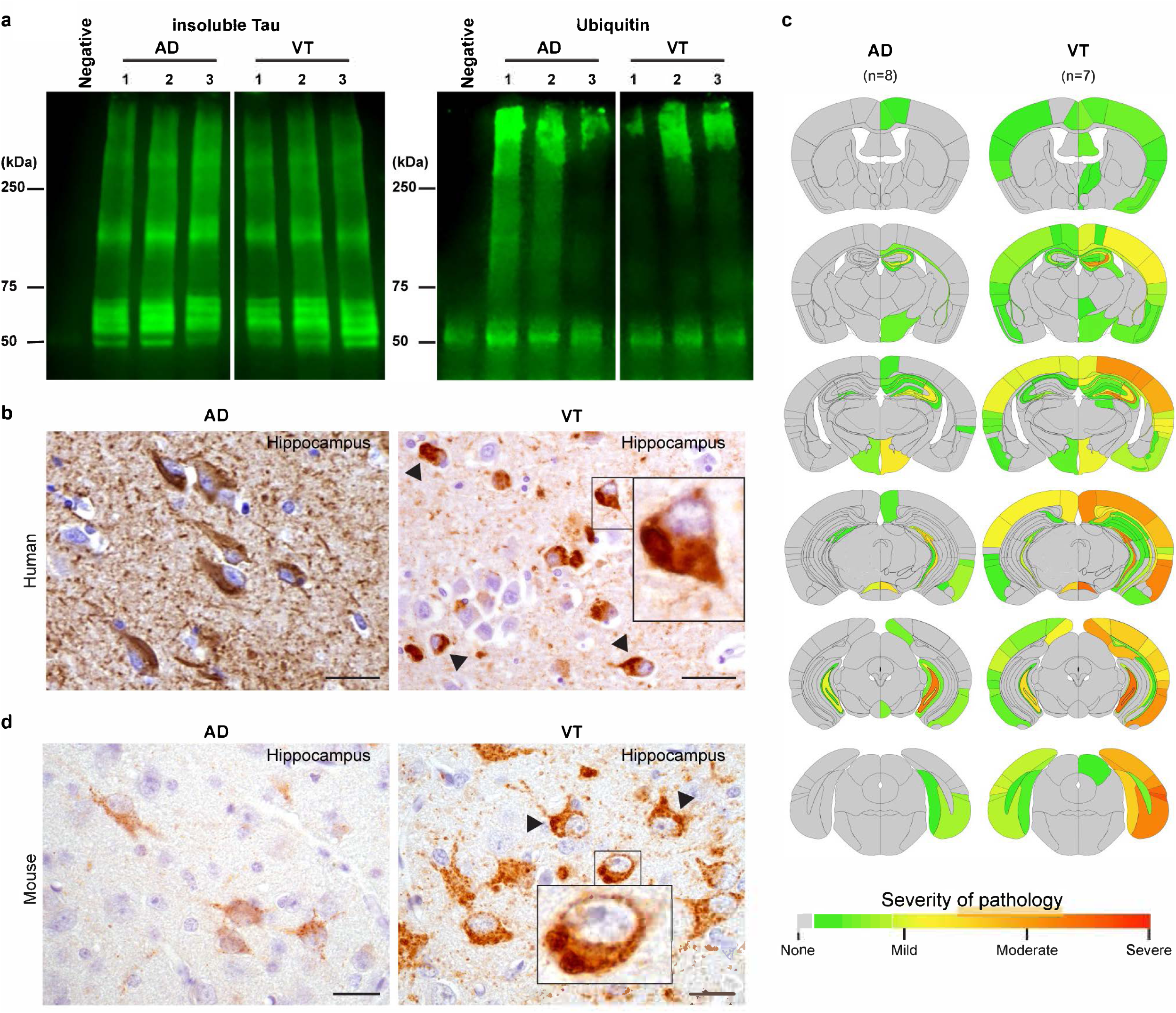
Intracerebral injection of WT mice with AD and VT brain-derived tau. (a) Ubiquitination of insoluble tau protein from AD and VT brains. Immunoprecipitation (IP) of sarkosyl-insoluble, tau-enriched brain lysates from AD and VT brains was performed using Tau-12 antibody and immunoblotted for either total tau protein (17025 rabbit antibody, left) or ubiquitin (E4I2J rabbit antibody, right). An IP without brain lysate (labeled “Negative”) was performed simultaneously to demonstrate a non-specific background band of ∼50 kDa. (b) Comparison of representative PHF-1-positive phosphorylated tau aggregates in the hippocampus of human brain tissue. AD (left, AD Case 2) shows many NFTs and neuropil threads, while VT (right, VT Case 1) shows neurofibrillary tangles in addition to neuronal cytoplasmic aggregates with a distinct, compact appearance (arrowheads and inset). (c) Semi-quantitative scoring of AT8-positive tau pathology in WT mice brains injected ipsilaterally (right side) with tau extracted from AD or VT brains. Pathology from 8 AD-injected and 7 VT-injected mice were scored from mild to severe (1-3, green to red), averaged, and mapped onto coronal mouse brain sections. Unaffected Regions are gray. (d) Comparison of representative AT8-positive tau aggregates in the hippocampus of injected WT mice brains. AD-injected mice (left) show diffuse, pretangle-like neuronal cytoplasmic staining, while VT-injected mice (right) show more intense and uneven neuronal cytoplasmic staining, including some round, compact aggregates (arrowheads and inset). Scale bars: (b) 40 μm, (d) 20 μm.

Despite these biochemical similarities between AD and VT tau, several neuropathological differences have been described. The distribution of tau pathology is different, with greater involvement of the medial temporal lobe including the hippocampus in AD compared to frontotemporal neocortex involvement in VT, resulting in distinct clinical phenotypes^11^. With respect to the morphology of the tau aggregates, VT brains exhibit neurofibrillary tangles in addition to occasional dense, round-shaped neuronal tau aggregates, in contrast with AD where such round inclusions are generally not observed (Fig. 1b).

To determine whether AD and VT tau filaments are associated with different biological activity *in vivo,* sarkosyl-insoluble tau from either AD or VT brain tissues were injected intracerebrally into 15 WT mice (n=8 for AD, 7 for VT). Mice were analyzed 3 months after injection by immunostaining brain sections with AT8, an antibody that recognizes phosphorylated tau. Blinded semiquantitative pathology scores of AT8-positive tau aggregates for each AD and VT injection were used to generate brain heat maps of the distribution and severity of tau pathology. AD tau-injected mice showed tau pathology mostly limited to the ipsilateral hippocampal regions, mimicking the preferential involvement of the medial temporal lobe in AD (Fig. 1c). In contrast, VT tau-injected mice showed more cortical and contralateral tau pathology, mimicking the preferential neocortical involvement seen in VT (Fig. 1c). In addition, AD tau-injected mice showed diffuse, pretangle-like neuronal cytoplasmic staining, while VT tau-injected mice showed more intense and uneven neuronal cytoplasmic staining including some compact, round inclusions (Fig. 1d). Thus, the distribution and morphology of tau pathology in injected mice brains with AD or VT tau appeared to recapitulate some features of each human disease.

### VT and AD tau filament structures

Recent structural studies have shown different tau filament core ultrastructures corresponding to different subcategories of tauopathy^1^. Different patterns of *in vivo* cell-to-cell transmission of tauopathy are hypothesized to be at least in part due to differences in tau filament core structure. Based on the distinct neuropathological features of AD and VT tau in both human and mouse brains, we hypothesized that, despite some biochemical similarities (i.e. tau isoform inclusion, insolubility, polyubiquitination, phosphorylation), AD and VT tau filament structures would be different. Sarkosyl-insoluble tau filaments were extracted from AD and VT tissues and analyzed using cryo-EM with helical reconstruction using RELION^13^. During the process of reconstruction, we employed several sequential 3D classification steps to successfully classify as many particles as possible (Extended Data Tables 3 and 4). As expected, reconstruction of tau filaments from two AD cases (same cases used for mouse injections above) showed the known paired helical filament (PHF) and straight filament (SF) cross-β sheet core structures consisting of two identical tau protofilaments^7^. Some additional densities adjacent to the sidechains of K317 and K321 were observed in averaged 2D cross-section density map images as previously reported^7^, particularly evident adjacent to the AD SF interface (Fig. 2a). The proportion of particle numbers contributing to AD PHF versus AD SF density maps was similar in both two cases where more than three-fourths of classified particles contributed to the AD PHF structure (Fig. 2b).

**Fig. 2.**
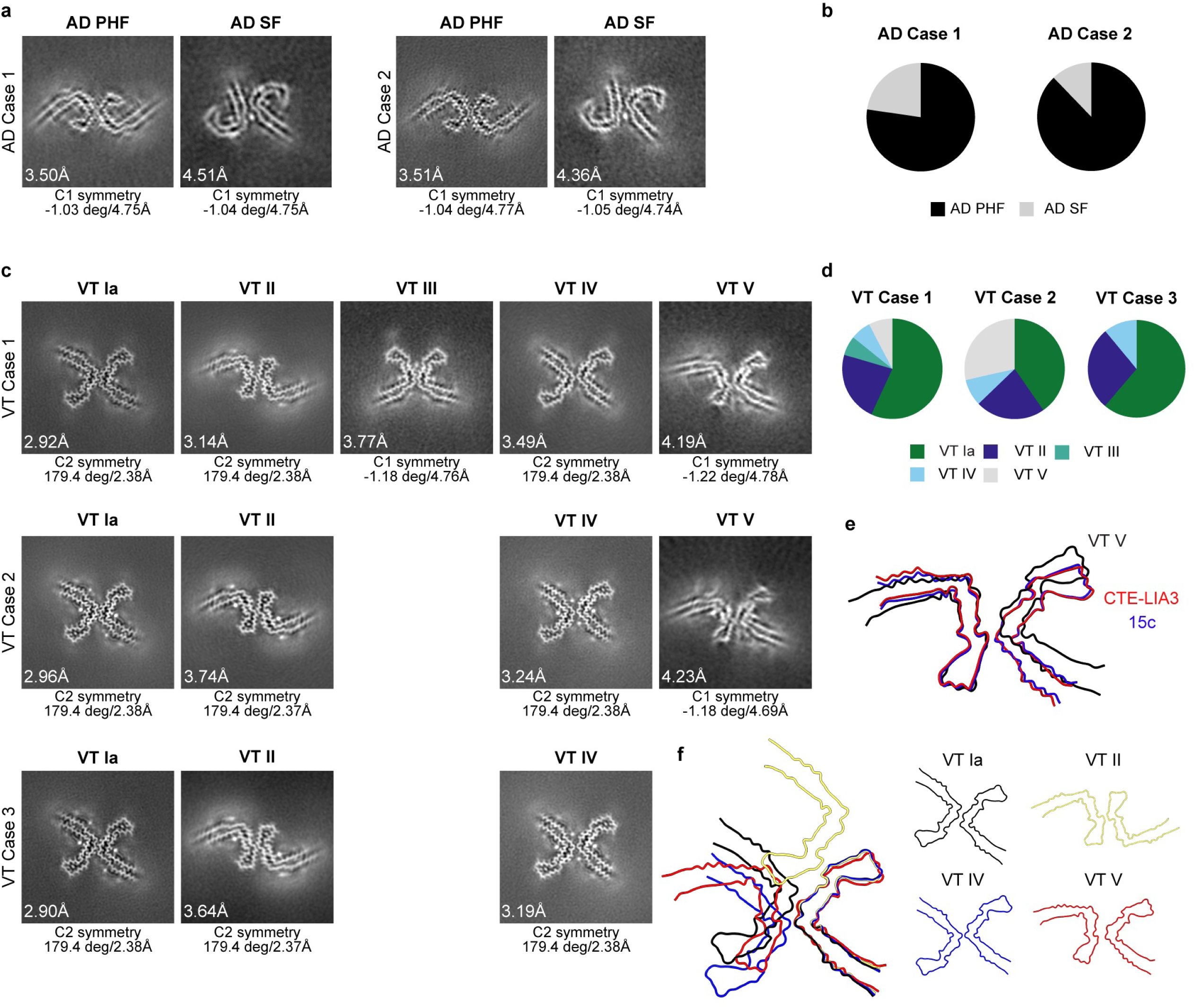
Reconstruction of AD and VT tau filament core structures. (a, c) 2D cross-sections of each refined density map of tau filament cores from two AD and three VT cases analyzed by cryo-EM. The resolution is shown in the lower left, with each filament’s symmetry, helical twist, and rise shown below each panel. (b, d) The proportion of particles contributing to the density map for each filament type with a resolution of 10 Å or higher. (e, f) Overlays of atomic models are shown to highlight different protofilament interfaces when comparing VT V (black), 15c (blue, PDB: 7QL0^14^), and CTE-LIA3 (red, PDB: 8Q9F^15^) cores (panel e) and VT Ia (black), VT II (yellow), VT IV (blue) and VT V (red) cores (panel f). Due to limited map resolution, the VT V model was made by manually overlaying VT tau protofilaments on the density map.

Tau filament reconstructions of the three VT cases resulted in five different cross-β sheet filaments, each consisting of the same pair of protofilaments but with differences in protofilament-protofilament interface interactions (Fig. 2c). Three of these VT cores correspond to those previously described in a single case of VT^6^, or with chronic traumatic encephalopathy (CTE) ^3^, amyotrophic lateral sclerosis/parkinsonism-dementia complex (ALS/PDC) of the island of Guam and the Kii peninsula of Japan^4^, and subacute sclerosing panencephalitis (SSPE) ^5^. We named these three VT filament types VT Ia, VT II, and VT III as they correspond to the known CTE type I through CTE type III cores^3–6^. Two additional VT filament types were identified which do not correspond to any known *in vivo* or *in vitro* generated tau filaments, designated VT IV and VT V, both exhibiting the same protofilament structure but with different protofilament-protofilament interfaces when compared to the other VT fibrils (Fig. 2c) ^3^. These five VT filaments were variably present across the three VT cases, with VT Ia, VT II, and VT IV filaments detected in all cases and VT III and VT V filaments seen in a subset of cases. Based on the proportion of particles contributing to each filament type, VT Ia was most abundant in all three cases, corresponding to 40.4% to 61.2% of particles, followed by VT II (22.4% to 27.7%) and VT IV (6.6% to 11.0%; Fig. 2d). AD filaments (PHF and SF) were not found in these three VT cases.

### AD and VT tau filament interfaces

The heterogeneity of VT filaments was associated with differences in their protofilament-protofilament interfaces. Thus, we compared VT filaments’ density maps and their corresponding atomic models to infer the factors which stabilize the protofilament interface of tau cores, including the presence/absence of 1) steric zippers, 2) hydrogen bonds, 3) electrostatic interactions between inversely charged residues, and 4) additional less-structured densities adjacent to the core filament structure (Fig. 2, 3, and Extended Data Fig. 1). VT Ia filaments had an ordered, approximate C2 symmetrical core structure where the protofilament interface forms a short steric zipper corresponding to residues S324 to H329 with two inter-protofilament hydrogen bonds (Fig. 2c and Extended Data Fig. 1b). VT Ia cores also exhibited additional densities adjacent to the core between residues L315-K317, K317-K321, and I360-K369 (Fig. 2c).

**Fig. 3.**
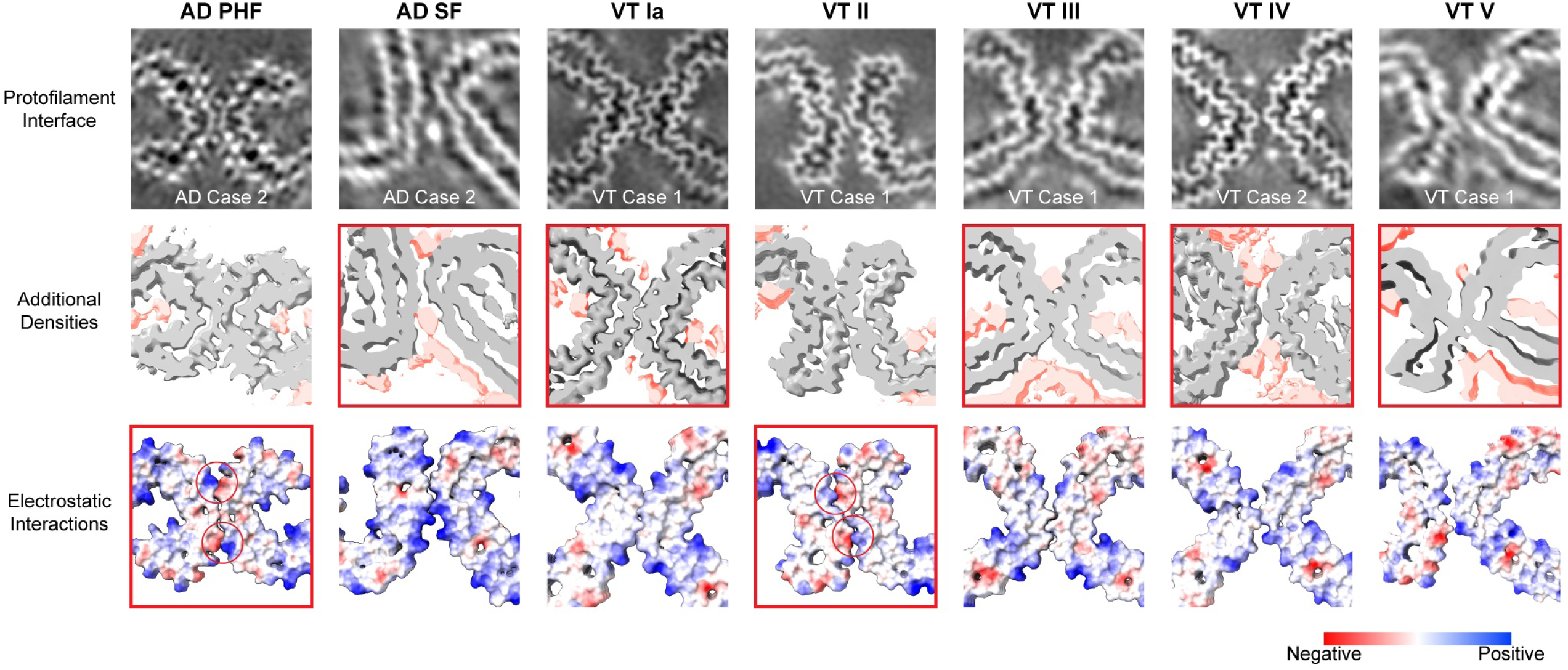
Structural characterization of AD and VT tau protofilament interfaces. The potential stabilizing factors contributing to each AD and VT filament core type are highlighted with red boxes. (Top) 2D cross-sections of the protofilament interface from each density map. The case number for each map is shown at the bottom of each panel. (Middle) 3D cross-sections corresponding to the density map in the top row are shown, corresponding to approximately three pairs of stacked protofilaments in the Z-axis. The contour level of each map is adjusted to show the additional densities next to the main chain, and these densities are highlighted in red. The panel is framed in red when clear additional densities are located adjacent to the protofilament interface. (Bottom) Atomic models are colored to show electrostatic potential (red and blue corresponding to negative and positive potentials, respectively). We used generated atomic models from the density map shown in the top panel except for AD PHF, AD SF, and VT III where published atomic models (PDB: 5O3O^7^, 5O3T^7^, and 8OT9^4^) are used. Due to limited map resolution, the VT V model was made by manually overlaying the VT tau protofilaments on the density map. The panel is framed in red if electrostatic inter-protofilament interactions exist between oppositely charged residues, with red circles indicating the interaction sites.

VT II had an ordered, approximate C2 symmetrical core structure with the protofilament interface formed by K331-E338 which included six inter-protofilament hydrogen bonds (Fig. 2c and Extended Data Fig. 1b). The VT II interface was further defined by two inter-protofilament interactions between the charged tau residues K331 and Q336/E338, which were not present in other VT protofilament interfaces (Fig. 3). The 2D cross-section of the VT II core showed additional densities associated with residues K317-K321 and I360-K369 which were distant from the interface (Fig. 2c).

The VT III filament had an asymmetric core structure with a protofilament interface consisting of a short steric zipper corresponding to residues G323-N327 and S324-I328 with five inter-protofilament hydrogen bonds (Fig. 2c and Extended Data Fig. 1b) ^4^. In contrast to the head-to-tail arrangement of protofilaments in all other VT fibrils, VT III was the only tau filament with a head-to-head arrangement. The VT III filament showed additional densities between the residues K317-K321 and I360-K369, in addition to H329 near the interface (Fig. 2c). In addition, a strong density between the paired K331 residues across the two protofilaments is reminiscent of the strong density that appears to stabilize the AD SF protofilament interface.

In contrast, the VT IV filament had an approximate C2 symmetrical core structure with a narrower protofilament interface formed by only two residues, G323 and S324, without a steric zipper structure and without any polar interactions (Fig. 3 and Extended Data Fig. 1). VT IV cores also exhibited additional densities adjacent to the core between residues L315-K317, K317-K321, and I360-K369 (Fig. 2c) where the extra densities between K317 and K321 that were immediately adjacent to the protofilament interface formed a relatively strong density in the 2D cross-section density map, similar to the strong interprotofilament densities that appears to stabilize AD SF and VT III cores.

The VT V filament had an asymmetric core structure with a protofilament interface formed by residues K321-S324 and G334-E338. While the resolution of the VT V core is relatively low, the VT V core also showed additional densities near the interface between residues E338-K340 and K317-T319 (Fig. 2c). We note that the VT V core has not been previously seen from human brain extracts, but is similar to *in vitro* core structures obtained from recombinant tau filaments (known as 15c^14^ and CTE-LIA3^15^). Fig. 2e shows an overlay of VT V filaments with 15c and CTE-LIA3. These two recombinant tau filament structures were characterized by electrostatic interactions between charged residues K321 and E338, which was less evident in the VT V filament^14,15^.

In summary, each VT filament core exhibited one or more factors which appear to stabilize the protofilament interface, perhaps driving the diversity of tau filament structures in disease. Interestingly, strong extra densities adjacent to the core appeared to stabilize AD SF, VT III, and VT IV cores. The VT III core was additionally stabilized by a steric zipper with multiple hydrogen bonds. In contrast, AD PHF and VT II protofilament interfaces did not exhibit strong extra densities but rather appeared to be stabilized by multiple hydrogen bonds in addition to electrostatic interactions between inversely charged residues at the protofilament interface. VT Ia cores were only stabilized by a short steric zipper with a few hydrogen bonds (Fig. 3 and Extended Data Fig. 1). To visualize the differences in protofilament positioning, an overlay of the four head-to-tail VT filament atomic models is shown (Fig. 2f).

### Shifting of VT tau filaments by ubistatin B

Given that both AD and VT tau are polyubiquitinated, we were interested in testing whether manipulating polyubiquitin can affect the ultrastructure of tau filament cores. To investigate this, we used a chemical approach, using ubiB to modulate the polarity of tau core polyubiquitin followed by visualization of tau filaments by cryo-EM^16^. UbiB is a small compound that can bind to two ubiquitins by hydrophobic and polar interactions. Structural models of a ubiquitin-ubiB-ubiquitin complex are presented in Fig. 4a^16^. Each ubiquitin’s surface is polar, and the surface of a ubiquitin-ubiB-ubiquitin complex shows an overall negative outer shell and relative shielding of positive residues internally thus altering the overall polarity of di-ubiquitins (Fig. 4a). We first validated that ubiB can bind to polyubiquitinated tau aggregates by incubating formalin-fixed AD and VT brain tissue sections with ubiB solution followed by immunolabeling sections with phospho-tau PHF-1 antibody. Confocal microscopy demonstrated colocalization of fluorescent ubiB to PHF1-positive tau aggregates in both AD and VT (Fig. 4b).

**Fig. 4.**
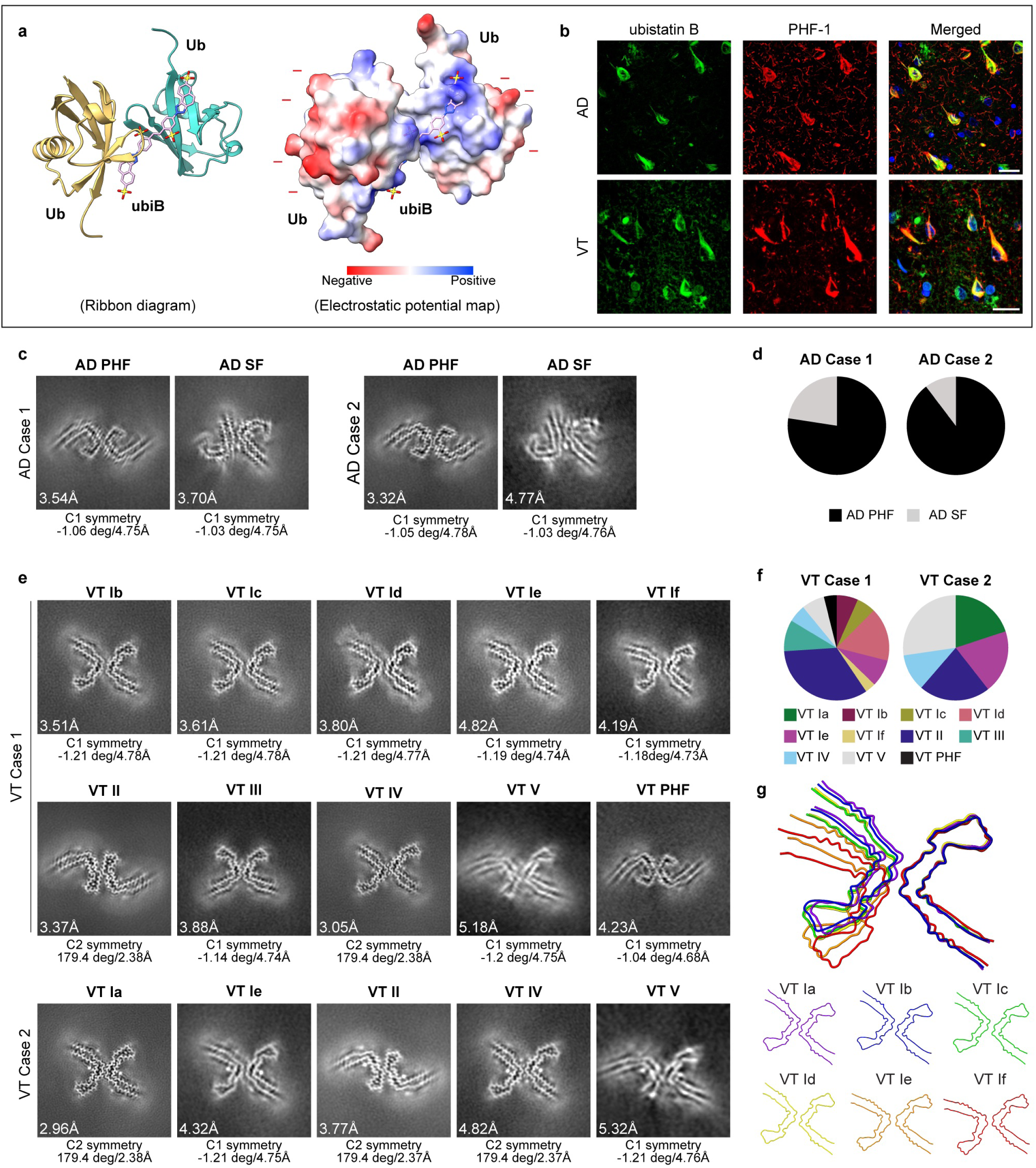
Ubistatin B-treated AD and VT tau filament core structures. (a) Structural models of the complex of di-ubiquitin and ubistatin B are shown as previously described^16^. The two ubiquitin molecules (PDB: 1UBQ^20^) are presented as either ribbon diagrams (left) or atomic surface models colored by electrostatic potential (right; red and blue show negative and positive potentials, respectively). Ubistatin B (PubChem SID: 475550170) bridges two ubiquitins, resulting in an overall negative outer shell and relative shielding of positive residues internally^16^. (b) Validation of the binding of ubiB to neurofibrillary tangles. Formalin-fixed, paraffin-embedded brain neocortex tissues from AD Case 2 and VT Case 2 were stained with 10 mM ubiB (green), followed by immunostaining with PHF-1 antibody (red) and a Draq-5 nuclear counterstain (blue). Scale bars: 20 μm. (c, e) 2D cross-sections of each refined density map of ubistatin B-treated tau filament cores from AD Case 1, AD Case 2, VT Case 1, and VT Case 2. The resolution is shown in the lower left, with each filament’s symmetry, helical twist, and rise shown below each panel. (d, f) The proportion of particles contributing to the density map for each filament type with a resolution of 10 Å or higher. (g) Overlays of our atomic models are shown to highlight shifting of protofilament interfaces from VT Ia (purple) to VT Ib (blue), VT Ic (green), VT Id (yellow), VT Ie (orange), and VT If (red) cores.

We then incubated tau filaments extracted from either AD Case 1, AD Case 2, VT Case 1, or VT Case 2 with ubiB solution, followed by cryo-EM reconstruction of tau filament core structures. The two ubiB-treated AD case samples showed the same AD PHF and AD SF filament types as detected in untreated preparations, including similar particle proportions corresponding to PHF (>75%) versus SF (<25%; Fig. 4c, 4d, and 5). In contrast, the reconstruction of ubiB-treated VT tau filaments yielded six filament types that were not detected in the untreated VT filament preparations (Fig. 4e, 4f, and 5). One of these six filaments corresponded to AD PHF (seen in VT Case 1, designated VT PHF in Fig. 4e). The other five new filaments had a superficially similar core morphology to that of VT Ia filaments. Hence, we categorized these filaments into the VT I group, designated VT Ib to VT If. All five of these filaments had an asymmetric core with slightly different protofilament interfaces formed by G323 to N327, each without the steric zipper seen in VT Ia. UbiB-treated tau filament preparations from VT Case 1 contained all five VT Ib through VT If types, while ubiB-treated tau filament preparations from VT Case 2 contained the VT Ie type. Interestingly, the VT Ia type was no longer detected in VT Case 1 and was reduced by 50.5% in VT Case 2 in ubiB-treated preparations (Fig. 4f). We generated atomic models from the density maps of VT Ib through If filaments and overlaid them with the atomic model for VT Ia to better compare their conformations (Fig. 4g and Supplementary Video. 1). Although these ubiB-dependent fibrils looked superficially similar, each had non-overlapping backbone structures with different types of inter-protofilament hydrogen bonds (Extended Data Fig. 1b). To ensure that the newly found VT Ib through If types do not exist within the untreated fibril datasets, the newly detected VT Ib through If density maps were used as reference maps for additional rounds of 3D classification for all unclassified particles in the untreated VT datasets. This failed to detect fibril types VT Ib through If in the untreated VT datasets.

Because ubiB alters the overall polarity of di-ubiquitin, we hypothesized that ubiB treatment shifted polyubiquitin moieties adjacent to the structured VT Ia core, thereby altering its protofilament-protofilament interface, resulting in VT Ib through VT If. To determine whether ubiB indeed shifted polyubiquitin, density maps of untreated filaments were compared to structures of ubiB-treated filaments. In general, upon ubiB treatment, some of the poorly structured additional densities adjacent to the structured core filament shifted towards electrostatically positively charged tau amino acid residues (colored blue in Fig. 6). Given the overall negative polarity of ubiB-treated di-ubiquitin (Fig. 4a), these shifting extra densities may represent polyubiquitin (Fig. 6). We visually present these shifting extra densities for AD PHF (Fig. 6a), AD SF (Fig. 6b), VT Ia (Fig. 6c), VT II, (Fig. 6d), VT III (Fig. 6e), and VT IV (Fig. 6f) filament cores, where “plus” signs correspond to the shifting of densities towards positively charged amino acids and “asterisks” correspond to regions where the density decreases upon ubiB treatment. Thick arrows correspond to the proposed shifting direction towards more positively charged residues.

In the ubiB-treated AD PHF, VT Ia, VT II, VT III, and VT IV density maps, densities around the positively charged K311, K317, and K321 residues were increased (Fig. 6a, 6c, 6d, 6e, and 6f). This corresponded to a decrease in density adjacent to the electrostatically neutral L315 in AD PHF, VT II, VT III, and VT IV density maps. The fact that there were areas of both increased and decreased density around the fibril core suggested that the shifting of densities are not due to differences in data quality or data processing artifacts. In AD PHF, VT II, and VT IV, a new density mass clearly emerged between K321-G323 (Fig. 6a, 6d, and 6f). AD SF showed increased density adjacent to K331. Densities towards the C-terminus of the structured core also increased around positively charged K343, R349, and between K375-R379 residues for ubiB-treated AD PHF, AD SF, VT Ia, VT II, and VT III. The shifting and concentration of these poorly structured extra densities towards positively charged residues upon ubiB treatment suggest that these represent polyubiquitin. In contrast, ubiB treatment did not change the relatively strong density present at the protofilament interface for AD SF between K317-K321 and VT III between K331 residues (Fig. 6b and 6e, thin arrows), and so their molecular identity remains unknown.

Finally, we returned to our examination of the protofilament interface of VT Ia through VT If filaments to understand how ubiB treatment resulted in the formation of these structures. For each of these filaments, including VT Ia, the inter-protofilament space between K321 and K331 residues was filled with increased additional density (Fig. 5, middle rows, and Fig. 6c). These densities at the protofilament interface were associated with widening of the inter-protofilament space between K321 and K331 for VT Ib through VT If compared to the VT Ia core based on both 2D cross-section density maps and atomic models (Fig. 5). Indeed, using our atomic models, the inter-protofilament distance between K321-K331 was greater for VT Ib (19.4 Å), VT Ic (21.5 Å), VT Id (20.8 Å), VT Ie (27.5 Å) and VT If (29.1 Å) than for VT Ia (17.4 Å) (Extended Data Fig. 2). Based on these findings, the accumulation of polyubiquitin around the positively charged lysine residues at the protofilament interface appeared to have pushed VT Ia protofilaments outwards, resulting in new protofilament interface structures stabilized by newly formed hydrogen bonds. Indeed, consecutive visualization of VT Ia through VT If appears to show that crowding of polyubiquitin towards the protofilament interface is associated with a processive outward shifting of protofilaments (Supplementary Video. 1).

**Fig. 5.**
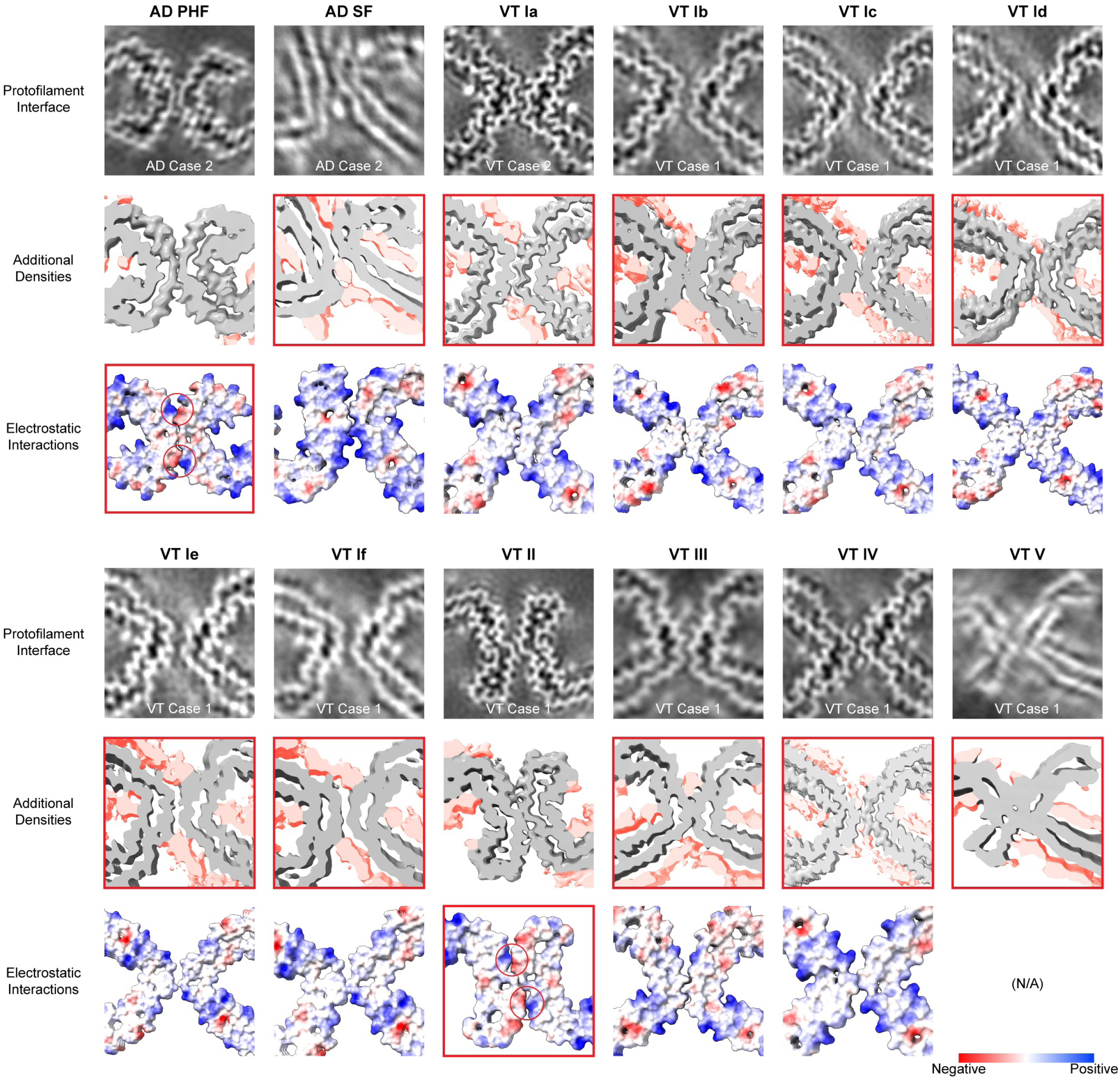
Structural characterization of ubistatin B-treated AD and VT tau protofilament interfaces. The potential stabilizing factors that contribute to each ubistatin B-treated AD and VT filament core type are highlighted with red boxes. (Top) 2D cross-sections of the protofilament interface from each density map. The case number for each map is shown at the bottom of each panel. (Middle) 3D cross-sections corresponding to the density map in the top row are shown, corresponding to approximately three pairs of stacked protofilaments in the Z-axis. The contour level of each map is adjusted to show the additional densities next to the main chain, and these densities are highlighted in red. The panel is framed in red when clear additional densities are located adjacent to the protofilament interface. (Bottom) Atomic models are colored to show electrostatic potential (red and blue corresponding to negative and positive potentials, respectively). Five atomic models (VT Ia, VT Ib, VT Ic, VT II, VT IV) were refined from our ubistatin B-treated datasets. The published atomic models (PDB: 5O3O^7^, 5O3T^7^, and 8OT9^4^) are used for AD PHF, AD SF, and VT III. Due to limited map resolution, the VT Id, VT Ie, and VT If models were made by manually overlaying protofilaments on the density map. The panel is framed in red if inter-protofilament interactions exist between the oppositely charged residues, with red circles indicating the interaction sites.

## Discussion

Various tau filament ultrastructures are related to the diversity of clinical and pathological phenotypes of tauopathies^1^. These tau filament structures have been hypothesized to be one factor that influences the cell-to-cell propagation of tau aggregates^17^, which in turn is thought to dictate the heterogeneous spatial distribution across different tauopathies.

We found that AD and VT tau have distinct biological activities *in vivo* upon inoculating WT mice with human brain tau extracts, mimicking their cognate human diseases in terms of both regional distribution of tauopathy and microscopic morphology of tau inclusions (Fig.1b, 1c, and 1d). This correlates with differences in the tau core filament structure determined by cryo-EM analysis of AD versus VT brain extracts. Some of this structural heterogeneity appears to be related to differences in protofilament interactions (Fig. 2 and 3). There are several structural features that appear to stabilize the protofilament interface, including electrostatic interactions, hydrogen bonds, and steric zippers (Fig. 3 and Extended Data Fig. 1). In addition, there are additional densities adjacent to filament core backbones which may also stabilize the protofilament interface (Fig. 3).

Similar additional densities have been observed in past structural reconstruction of pathologic tau filament cores and have been postulated to be PTMs or other unknown cofactors^3,7^. Recent studies combining proteomics and cryo-EM methods have found disease-specific patterns of tau filament core PTMs between different tauopathies^10^, suggesting that PTMs including ubiquitination contribute to disease heterogeneity in part by stabilizing inter-protofilament packing. We have inferred the location of ubiquitin PTMs on the tau filament core by chemically stabilizing and shifting polyubiquitin chains with ubiB (Fig. 6). The shifting of VT tau core ubiquitins caused multiple new core protofilament conformations, demonstrating that a subset of tau filaments exhibit some level of fibril instability where PTM localization can influence the protofilament-protofilament interface. These results suggest that structural elements outside of the highly ordered tau filament core, including PTMs, may influence filament structure and stability.

**Fig. 6.**
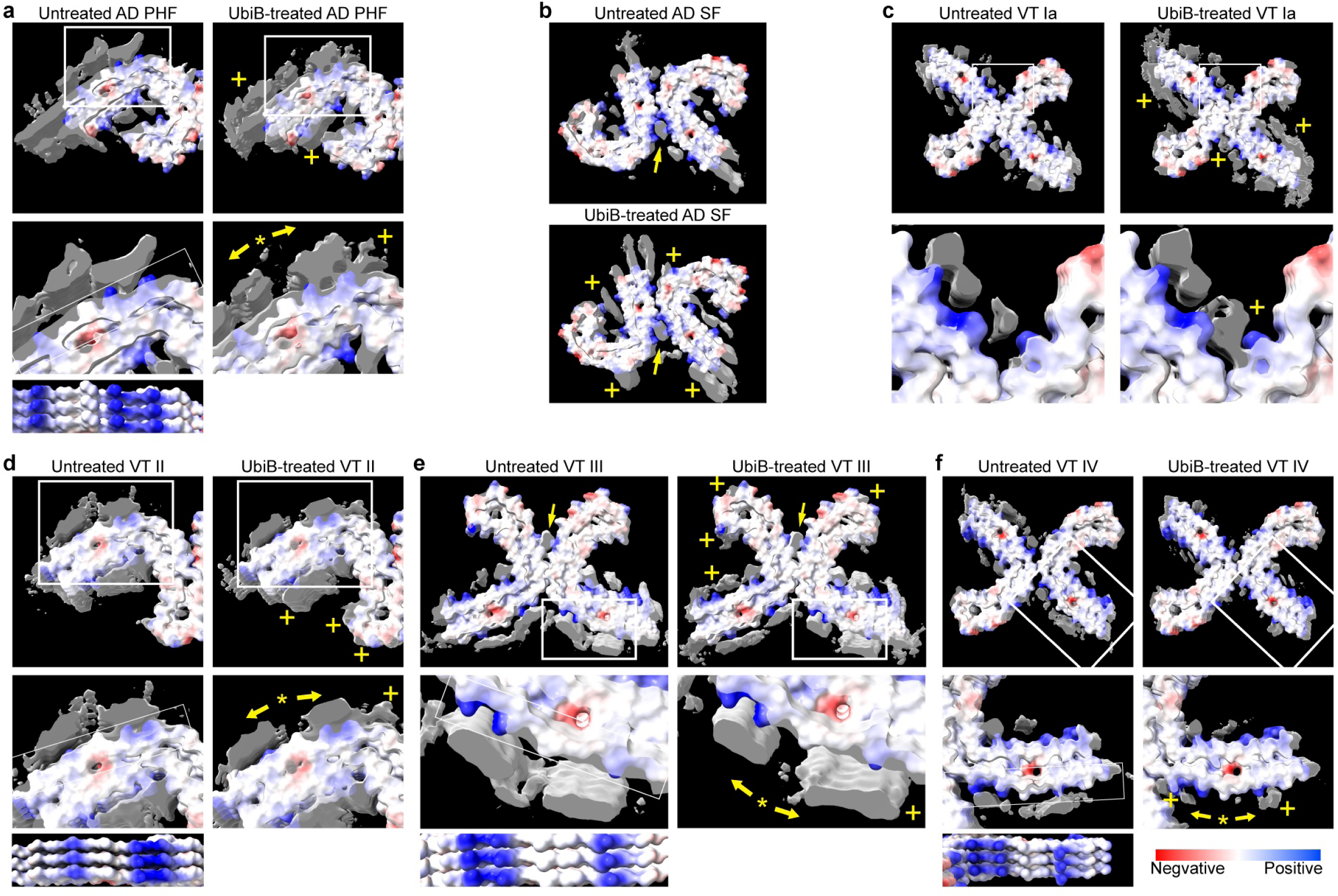
Shifting additional densities within ubistatin B-treated AD and VT tau filament cores. Additional densities adjacent to the untreated and ubiB-treated tau filament cores are compared for (a) AD PHF, (b) AD SF, (c) VT Ia, (d) VT II, (e) VT III, and (f) VT IV. Density map and refined atomic model overlays are shown with core surfaces colored based on electrostatic potential (red and blue corresponding to negative and positive potentials, respectively). Panels a, d, e, and f additionally contain slightly different views of the atomic model to better visualize the tau filament surface electrostatic potentials. Each density map corresponds to a thickness of approximately three pairs of stacked protofilaments in the Z-axis. Each map’s contour level is adjusted such that the structured core filament is approximately the same width when comparing untreated and ubiB-treated maps. The shifted densities in ubiB-treated maps are indicated with a “plus” for an increase, an asterisk for a decrease, and thick arrows corresponding to the proposed shifting direction towards more positively charged residues. Thin arrows in panels b and e highlight additional densities immediately adjacent to the protofilament interface which did not shift upon ubiB treatment. The representative density maps were taken from AD Case 2 (a, b), VT Case 1 (d-f), and VT Case 2 (c). We used published atomic models (PDB: 5O3O^7^, 5O3T^7^, and 8OT9^4^) (a, b, e) and our atomic models from VT Case 1 (d, f) or VT Case 2 (c).

In this study, the analyzed tau filaments were extracted and isolated from diseased human brain tissue. Thus, their native distributional and interatomic features were not available. In the future, visualizing these tau ultrastructures *in situ* using cryo-electron tomography methods recently developed for human brain tissue^18,19^ could provide additional insights into the pathophysiology of tauopathies, including the cell-to-cell transmission of protein aggregates, to better understand the molecular mechanisms contributing to clinical and pathological heterogeneity. In addition, cryo-EM helical reconstruction has provided static images of VT tau fibrils in different conformations. Given the structural lability of the VT Ia protofilament interface, which was notably not seen for AD tau filaments, there is the possibility that there is actually a dynamic range of shifting tau filaments *in vivo*, some of which may be influenced by PTMs or other intracellular factors.

In conclusion, a combined functional, structural and chemical biology approach has allowed for the localization of polyubiquitin adjacent to tau filament cores and has provided evidence that features outside of the highly structured tau core can contribute to ultrastructural heterogeneity.

## Methods

### Human autopsy brain tissues

We determined the core ultrastructure of tau filaments from the brains of five autopsied cases: AD Case 1, AD Case 2, VT Case 1, VT Case 2, and VT Case 3. Autopsied human brain tissue was obtained from the University of Pennsylvania Center for Neurodegenerative Disease Research Neurodegenerative Disease Brain Bank, where all cases were diagnosed pathologically as AD or VT^11^. Informed consent for autopsy was obtained from next of kin for all cases. Demographic and pathological information is available in Extended Data Table 1 or our previous VT case series publication^11^.

### Mice

C57BL/6J female mice were obtained from the Jackson Laboratory and maintained in a pathogen-free facility at the University of Pennsylvania. All experimental protocols were approved by the Institutional Animal Care and Use Committee (IACUC) of the University of Pennsylvania.

### Extraction of sarkosyl-insoluble tau

AD and VT tau filaments were sequentially extracted from frozen, human frontal neocortex as previously described^8^ with minor modifications. All extracted cases were screened to exclude cases with comorbid neocortical Lewy body and TDP-43 pathologies by immunohistochemical examination. ∼13 grams of grey matter were separated from white matter using a scalpel. Brain homogenate was prepared in nine volumes (v/w, ml/g) of PHF buffer (10 mM Tris, pH 7.4, 10% sucrose, 0.8 M NaCl, 1 mM EDTA) with 0.1% sarkosyl, 0.1 mM PMSF, 2mM DTT, proteinase and phosphatase inhibitor cocktails (Thermo Scientific) in a glass dounce homogenizer and spun at 10,000 xg for 10 min at 4 °C. The supernatant (sup 1) was collected, and the pellet was extracted again using the same volume of PHF buffer. The supernatant (sup 2) and sup 1 were then pooled together and filtered, and 25% sarkosyl solution was added to achieve a final concentration of 1% (total sup). Total sup was incubated in a beaker with stirring for 60-90 min at RT and followed by a 125,400 xg spin for 75 min at 4 °C. The pellet was collected, washed twice with PBS, and resuspended in PBS with mild sonication (sark pel) of which a small portion was saved for cryo-EM. The rest of the sark pel was spun at 120,000 xg for 30 min at 4 °C, and the pellet was resuspended in PBS, extensively sonicated, and spun at 10,000 xg for 10 min at 4 °C. The tau-enriched supernatant (final sup) was collected and stored at −80 °C for immunoprecipitation and mice experiments.

### Quantification of protein concentrations

The total protein and tau concentrations in the extracted sark pel and final sup lysates were determined using a BCA assay kit (Thermo Scientific) and quantitative immunoblot, respectively. The quantified results are shown in Extended Data Table 2. For immunoblot, each lysate was diluted x50 fold in Laemmli Sample Buffer, run on a 4-20% gradient sodium dodecyl sulfate-polyacrylamide gel electrophoresis (SDS-PAGE) gel, and transferred to nitrocellulose membranes. Membranes were then blocked in MB-070 blocking buffer (Rockland) at RT for 1 hour and immunoblotted with specific primary antibodies (Extended Data Table 7). The blots were further incubated with IRDye-labeled secondary antibody (LI-COR) and visualized using an ODYSSEY Fc imager (LI-COR). The bands were quantified using ImageJ software (National Institutes of Health) ^21^.

### Comparison of the ubiquitination levels within immunoprecipitated insoluble tau from AD and VT Brains

Tau immunoprecipitation was performed using Tau-12 antibody (monoclonal, Biolegend) and Dynabeads Protein G Immunoprecipitation Kit (Invitrogen) according to a previous publication^12^ and manufacturer’s instructions with modifications. 900 μg of Dynabeads were resuspended by rotating for 5 min. The supernatant was removed using a magnetic stand, and beads were washed in PBS+0.05% Tween20. To reduce the immunoblot bands derived from bead-bound Protein G, beads were resuspended in 30 μL of Laemmli Sample Buffer, incubated twice at 95 °C for 5 min, and washed in PBS+0.05% Tween20. Beads were resuspended in 100 μL of PBS containing 5 μg of Tau-12 and incubated for 20 min at RT with rotation. The supernatant was removed using a magnetic stand, and bead-antibody complexes were washed in PBS+0.05% Tween20. The tau-enriched, extensively sonicated lysates extracted from each of the three AD and three VT brains were diluted with SDS in PBS solution to achieve a final concentration of 2% SDS which were subsequently incubated at RT for 1 hour to denature tau protein. Denatured tau preparations were diluted with PBS to achieve a final volume of 200 μL corresponding to 0.1% SDS. The denatured tau lysates were added to the bead-antibody complexes and incubated for 2 hours at RT with rotation, and the supernatant was removed using a magnetic stand. After washing twice in PBS+0.05% Tween20 and then in PBS, bead-antibody-antigen complexes were resuspended in 15 μL of Laemmli Sample Buffer and incubated for 5 min at 95 °C. The supernatant was collected using a magnetic stand for immunoblotting using antibodies listed in Extended Data Table 7.

### Mice stereotaxic surgery and tissue processing

Tau-enriched final sup lysate from AD Case 1, AD Case 2, VT Case 2, or VT Case 3 were used for the injection experiments on eight and seven mice at 8-12 weeks of age for AD and VT injections, respectively. Mice were anesthetized with ketamine-xylazine-acepromazine, placed in a stereotaxic frame (David Kopf Instruments), and aseptically injected with the lysate in the right dorsal hippocampus and overlying cortex (bregma: −2.5mm, lateral: +2mm, depth: −2.4 mm and −1.4 mm from skull) with a Hamilton syringe. Each injection site received 2.5 μL of inoculum at 0.4 μg tau/μL as previously described^8^. Mice were anesthetized 3 months after injection and then sacrificed by transcardiac perfusion with PBS. The whole brain was dissected and fixed overnight in formalin for immunohistochemical analyses.

### Immunohistochemistry

Six μm formalin-fixed, paraffin-embedded human and mice tissue sections were stained using immunohistochemistry or immunofluorescence as previously described^22^ using antibodies listed in Extended Data Table 7. Blinded semiquantitative analysis of tau pathology was performed for mice brains by scoring the neuronal tau pathology in designated regions in coronal tissue sections. The scoring was performed manually by evaluating AT8-positive tau aggregates as 0-3 (0=no pathology, 3=severe pathology) for each mouse. Averaged scores from AD and VT-injected groups were used to generate heat maps representing the severity and distribution of tau pathology.

### Cryo-EM

Extracted sark pel lysates were diluted in PBS and briefly spun at 3,000 xg for 1 min. 3 μL of the supernatant was applied to glow-discharged holey carbon grids (QuantiFoil Cu R1.2/1.3, 200 mesh) and plunge-frozen in liquid ethane using a Leica EM GP2 Automatic Plunge Freezer. Images were acquired using EPU software on a Gatan K3 Summit detector in counted super-resolution mode using a Thermo Fisher Titan Krios G3i at 300 kV. The Gatan energy filter was used with a slit width of 20 eV. Further details are given in Extended Data Tables 5 and 6.

### Ubistatin B staining of formalin-fixed brain tissue sections

UbiB was synthesized following previously published procedures^16^ and was diluted in PBS. To validate the specific binding of ubiB to the polyubiquitinated tau filaments, six μm formalin-fixed, paraffin-embedded AD or VT frontal cortex sections were deparaffinized, hydrated and incubated with 10 μM ubiB in PBS at RT for 6 hours in the dark. The sections were then blocked with 2% FBS in 0.1M Tris buffer, pH 7.6, and immunostained overnight with PHF-1 mouse antibody, which was visualized with an anti-mouse fluorescently labeled secondary antibody (Invitrogen Alexa Fluor 568). Nuclear counterstain was done with Draq-5 (Cell Signaling Technology). Fluorescent images were obtained by a Leica TCS SPE laser scanning confocal microscope, using 405 nm, 561 nm, or 635 nm excitation wavelengths to detect ubiB, PHF-1, or Draq-5, respectively.

### Ubistatin B treatment of tau filaments for cryo-EM

The sark pel lysates from AD Case 1, AD Case 2, VT Case 1, and VT Case 2 were briefly spun at 3,000 xg for 1 min, and the supernatant was diluted in ubiB solution to achieve a mixture of 1 mM ubiB and ∼2.5 μM tau protein in PBS. The mixture was incubated at RT for 4 hours in the dark, and 3 μL of the mixture was applied on a carbon grid for cryo-EM as described above.

### Helical reconstruction

Multi-frame super-resolution movies (0.54 Å/pixel) were corrected for gain reference, binned by a factor of 2 (1.08 Å/pixel), motion-corrected, dose-weighted, and then summed into a single micrograph using the motion correction program of RELION-3.1.4 or RELION 5^23^. Aligned, non-dose-weighted micrographs were used to estimate the contrast transfer function (CTF) using CTFFIND-4.1^24^. All subsequent image-processing steps were performed using helical reconstruction methods in RELION-3.1.4 or RELION 5^13,25^. Filaments were picked manually in the micrographs for all datasets, and particles were extracted using a box size of 384 pixels and an inter-box distance of 9.4 Å. The reference-free 2D classification was performed using a regularization value of T = 2, and particles in suboptimal 2D classes were discarded. The remaining particles were used for the initial 3D classification using a featureless cylinder with a 384-pixel diameter as a reference map, with an estimated rise of 4.75 Å, and −1- and −1.2-degree of helical twists for AD and VT cases, respectively, based on the observed cross-over distances of filaments in the micrographs. Then, the sequential 3D subclassifications and 3D auto-refinements were combined to select the best density map for each structure. In these 3D classifications and refinements, a reference map was employed from the preceding runs of the same dataset except for the following: for reconstructing AD SF cores in two AD cases, a 3D subclassification was done for the subset of particles after removing the particles contributing to the AD PHF type, using a published density map for AD SF (EMD-3743) ^7^ as the reference with the 40 Å initial low-pass filter; for reconstructing ubiB-treated tau filaments, a density map of the untreated tau filament from the same case was employed as the reference with the 40 Å initial low-pass filter. The density maps with the best resolution for each filament type were then subject to Bayesian polishing and CTF refinement to improve the resolution further^26^. Final reconstructions were sharpened, and final, overall resolution estimates were calculated from Fourier shell correlations at 0.143 between the two independently refined half-maps, using phase-randomization to correct for convolution effects of a generous, soft-edged solvent mask using the post-processing procedures in RELION-3.1.4 or RELION 5^27^. Subsequent to the reconstruction for all untreated and ubiB-treated VT case datasets, additional 3D classifications were attempted for all unclassified particles in both untreated VT datasets using the newly detected VT I group density maps from the same VT case’s ubiB-treated dataset as a reference using a 40 Å initial low-pass filter. Figures were prepared using ChimeraX^28^. The XY cross-sectional images of the density map were generated by averaging the densities in four successive Z-sections corresponding to approximately 4.3 Å. The number of particles contributing to the density maps for each filament type with a resolution greater than 10 Å was summed to calculate the proportion of each filament type. Further details are described in Extended Data Figs. 1-3 and Extended Data Tables 3-6.

### Atomic model building and refinement

The atomic models were generated for the untreated VT Ia, VT II, and VT IV filaments, and the ubiB-treated VT Ia, VT II, VT Ib, VT Ic, and VT IV filaments from each density map selected by the best resolution from our datasets. The initial model was made by automatic model generation in ModelAngelo^29^ and manual alignments using ChimeraX^28^ and COOT^30^. The nine adjacent cross-β rungs from each initial model were retained for the real-space refinements using COOT^30^ and PHENIX^31^. Figures were prepared using ChimeraX^28^. Further details of the above are described in Extended Data Tables 5 and 6. We manually aligned our VT-fold tau protofilaments on VT Id, VT Ie, VT If, and VT V density maps to generate line models of the fibril backbone or for atomic surface models, but we did not generate a refined atomic model for these four types due to their relatively lower map resolutions.

## Acknowledgments

This study was supported by grants from the NIH (RF1AG065341, P30AG072979, and P01AG066597 to E.B.L.; and R35GM142505 to G.M.B.), the DeCrane Family Fund for PPA Research, and by gifts from the Shanahan Family Foundation and the Barrist Family Foundation. We thank Ruchao Peng and Longsheng Lai from the Yi-Wei Chang Lab and Shafiqur Rahman from the George Burslem Lab at the University of Pennsylvania, Stefan Steimle and Prerana Gogoi from the Beckman Center for Cryo-Electron Microscopy, and the Center for Neurodegenerative Disease at the University of Pennsylvania for their support and technical assistance. We thank our patients and their families who made this study possible and for their perseverance.

## Contributions

R.W., N.F.D., and E.B.L. examined human neuropathology; R.W. and L.C. performed biochemical analyses; R.W., N.F.D., H.X., K.H., B.Z., and C.M.O. performed animal experiments; R.W. and B.C.C. performed cryo-EM; R.W., B.C.C., Y.C., and E.B.L. analyzed cryo-EM data; R.W. and B.C.C. built atomic models; V.M.L. provided oversight for biochemical analyses and animal studies; G.M.B. and E.B.L. obtained the funding; E.B.L. supervised the project; all authors contributed to writing and review of the manuscript.

## Additional information

Supplementary information is available for this paper. Correspondence and requests for materials should be addressed to Edward B. Lee.

## Data availability

The cryo-EM map data have been deposited to the Electron Microscopy Data Bank (EMDB) under accession numbers: EMD-49670 for AD PHF and EMD-49671 for AD SF (AD Case 1); EMD-49672 for AD PHF and EMD-49675 for AD SF (AD Case 2); EMD-49627 for VT Ia, EMD-49629 for VT II, EMD-49678 for VT III, EMD-49680 for VT IV, and EMD-49681 for VT V (VT Case 1); EMD-49682 for VT Ia, EMD-49683 for VT II, EMD-49638 for VT IV, and EMD-49684 for VT V (VT Case 2); EMD-49686 for VT Ia, EMD-49689 for VT II, and EMD-49691 for VT IV (VT Case 3); EMD-49693 for AD PHF and EMD-49694 for AD SF (ubiB-treated AD Case 1); EMD-49695 for AD PHF and EMD-49697 for AD SF (ubiB-treated AD Case 2); EMD-49647 for VT Ib, EMD-49648 for VT Ic, EMD-49698 for VT Id, EMD-49699 for VT Ie, EMD-49700 for VT If, EMD-49650 for VT II, EMD-49701 for VT III, EMD-49658 for VT IV, EMD-49703 for VT V, and EMD-49702 for VT PHF (ubiB-treated VT Case 1); EMD-49641 for VT Ia, EMD-49704 for VT Ie, EMD-49705 for VT II, EMD-49706 for VT IV, and EMD-49707 for VT V (ubiB-treated VT Case 2). Refined atomic models have been deposited to the Protein Data Bank (PDB) under accession numbers 9NPK for VT Ia (VT Case 1), 9NQ4 for VT Ia (ubiB-treated VT Case 2), 9NQB for VT Ib (ubiB-treated VT Case 1), 9NQC for VT Ic (ubiB-treated VT Case 1), 9NPQ for VT II (VT Case 1), 9NQE for VT II (ubiB-treated VT Case 1) 9NQ0 for VT IV (VT Case 2), and 9NQI for VT IV (ubiB-treated VT Case 1).

## Extended Data Figures and Tables

**Extended Data Fig. 1.**
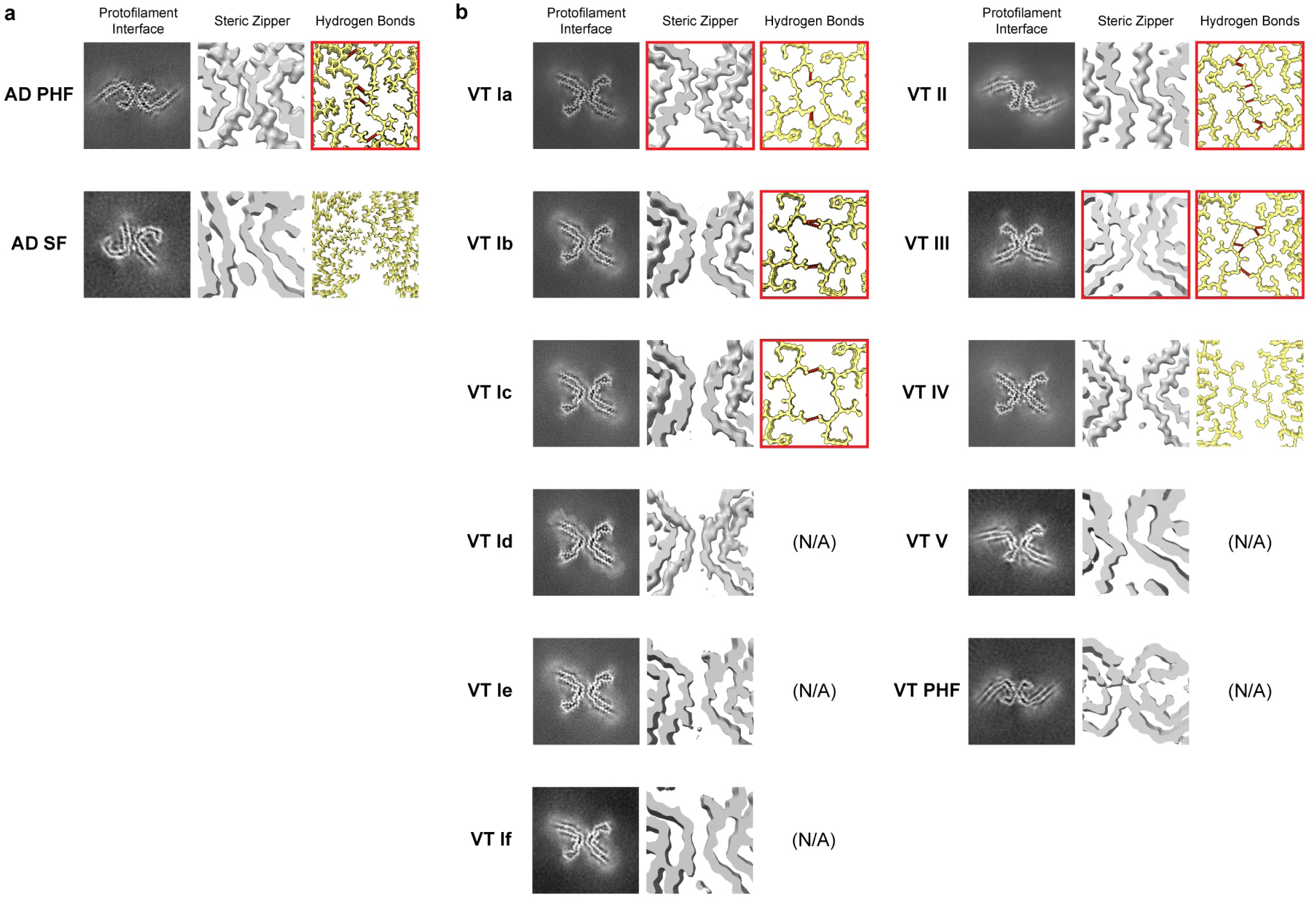
Steric zipper and hydrogen bonds within AD and VT tau protofilament interface. Comparison of inter-protofilament steric zipper and hydrogen bond features that appear to stabilize the protofilament interface. For each presented AD type (a) and VT type (b) tau filament core, the left and middle panels show the 2D and 3D cross-sections from density maps, respectively. The contour levels of 3D cross-sections are adjusted to emphasize the backbone chain structure. The right panels show the atomic model for each tau filament as a stack of three consecutive monomers. Inter-protofilament hydrogen bonds are visualized as red dotted lines using ChimeraX. The published atomic models (PDB: 5O3O^7^, 5O3T^7^, and 8OT9^4^) are used for AD PHF, AD SF, and VT III. We do not present the hydrogen bond positions for VT Id, VT Ie, VT If, VT V, and VT PHF types since we could not generate their refined atomic model due to limited map resolutions.

**Extended Data Fig. 2.**
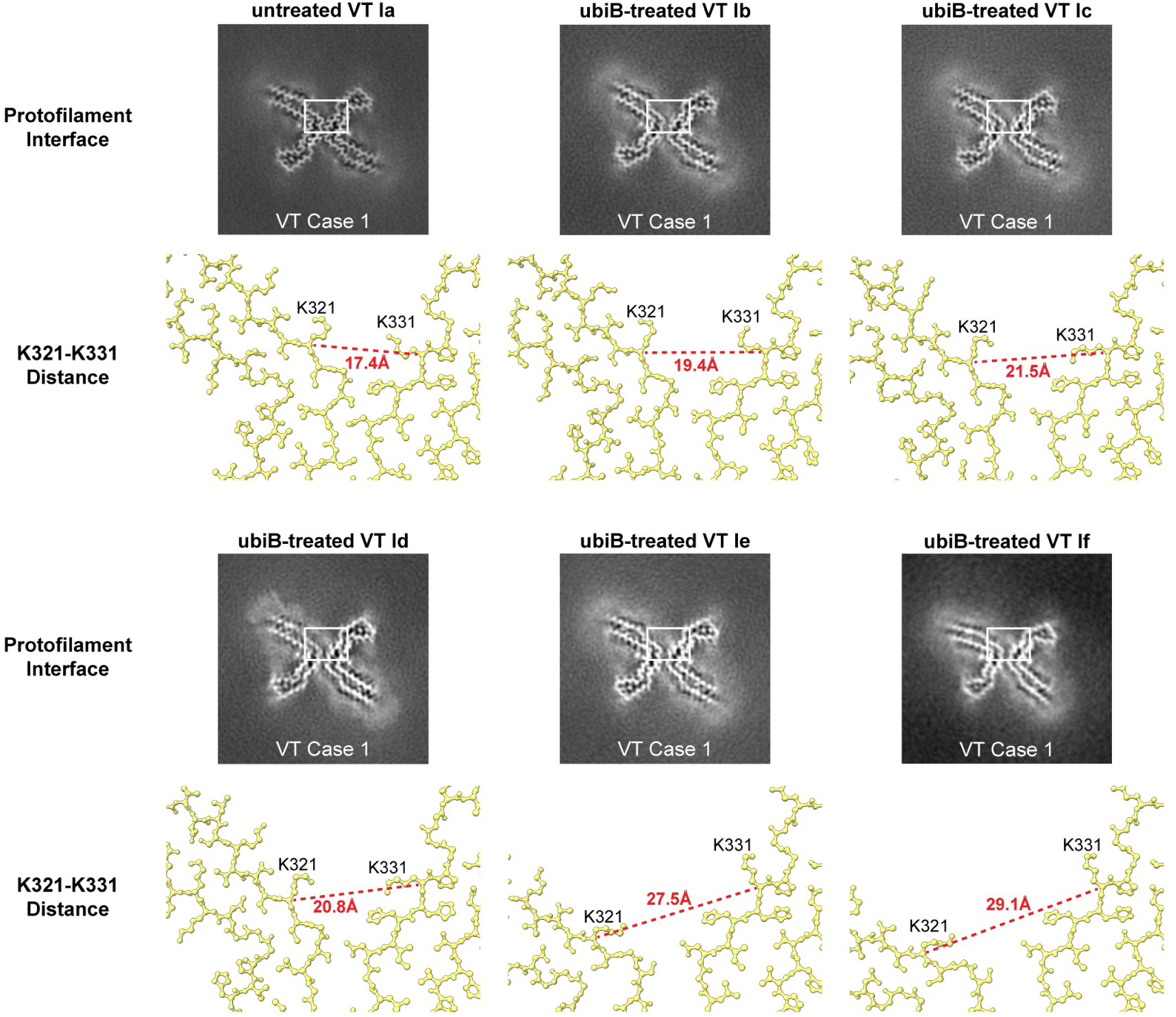
The distance between inter-protofilament K321-K331 residues in VT I tau filaments. A portion of each atomic model of untreated VT Ia filaments and ubiB-treated VT Ib, VT Ic, VT Id, VT Ie, and VT If filaments are shown, corresponding to the rectangular areas in the 2D cross-section image above. The distances between the central (α-) carbon atoms of K321 and K331 amino acids that span the inter-protofilament space are shown in red. The same scale is used for each panel to demonstrate the differences in distance.

**Extended Data Fig. 3.**
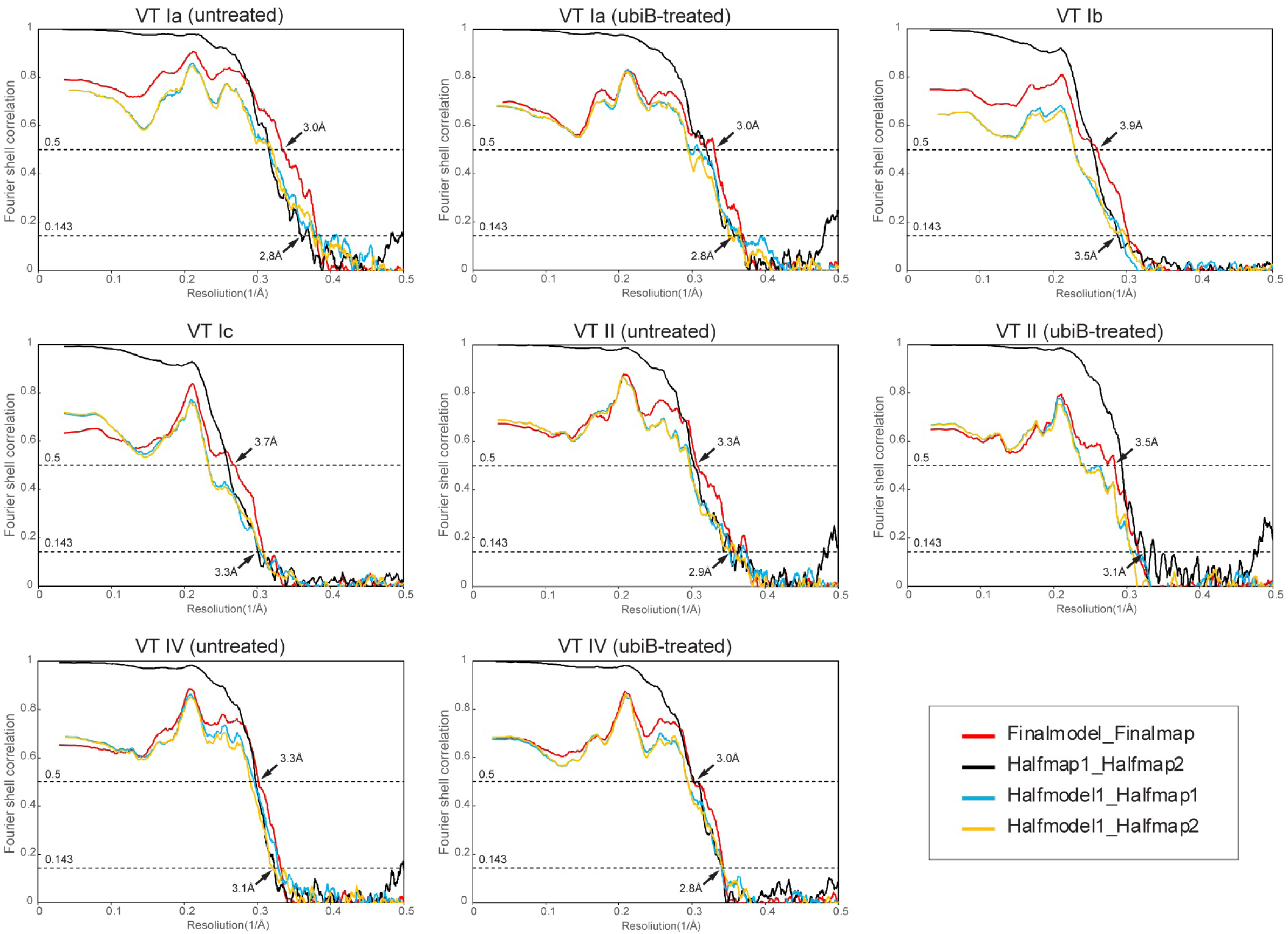
Fourier shell correlation (FSC) curves for cryo-EM maps and atomic models. Fourier Shell Correlation (FSC) curves for density maps and atomic models of VT filaments are shown for VT Ia (VT Case 1); ubiB-treated VT Ia (VT Case 2); VT Ib (VT Case 1); VT Ic (VT Case 1); VT II (VT Case 1); ubiB-treated VT II (VT Case 1); VT IV (VT Case 2); and ubiB-treated VT IV (VT Case 1). FSC curves correspond to two independently refined half-maps (half-map 1 vs. half-map 2) (black); the final refined atomic model against the final map (red); the atomic model generated on half-map 1 against half-map 1 (blue); and the atomic model generated on half-map 1 against half-map 2 (yellow).

**Extended Data Table 1.**
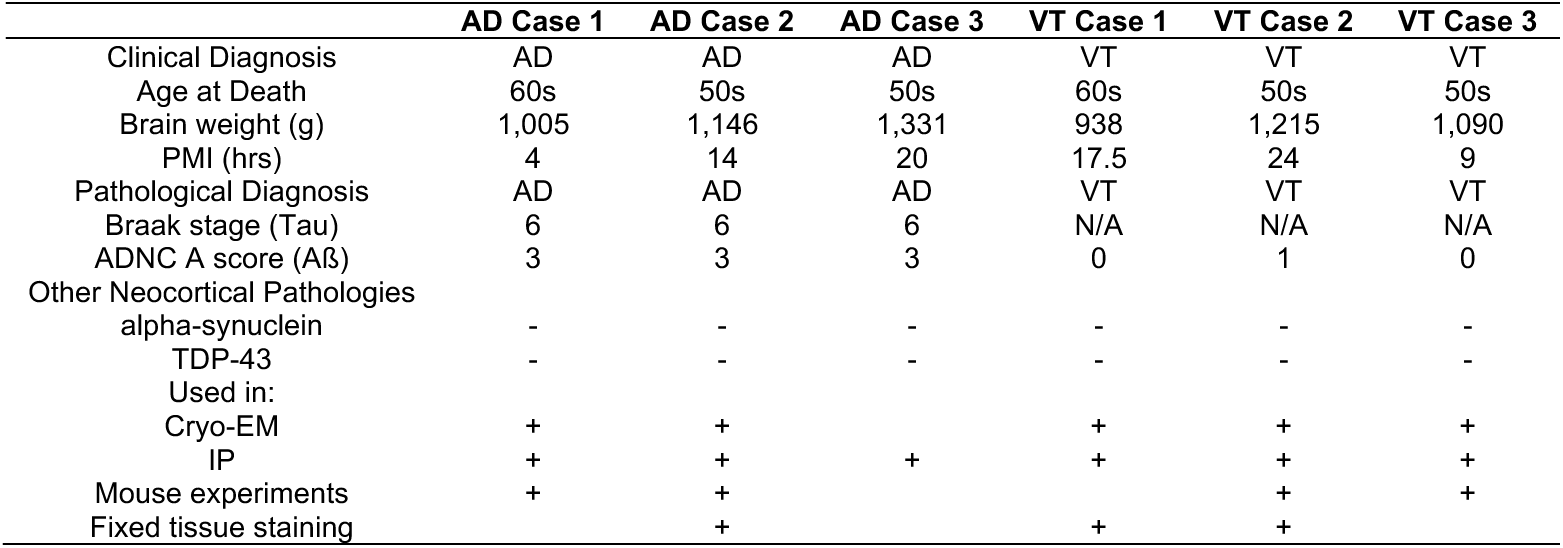
Demographic and pathological characteristics of AD and VT cases.

|  | AD Case 1 | AD Case 2 | AD Case 3 | VT Case 1 | VT Case 2 | VT Case 3 |
| --- | --- | --- | --- | --- | --- | --- |
| Clinical Diagnosis | AD | AD | AD | VT | VT | VT |
| Age at Death | 60s | 50s | 50s | 60s | 50s | 50s |
| Brain weight (g) | 1,005 | 1,146 | 1,331 | 938 | 1,215 | 1,090 |
| PMI (hrs) | 4 | 14 | 20 | 17.5 | 24 | 9 |
| Pathological Diagnosis | AD | AD | AD | VT | VT | VT |
| Braak stage (Tau) | 6 | 6 | 6 | N/A | N/A | N/A |
| ADNC A score (Aβ) | 3 | 3 | 3 | 0 | 1 | 0 |
| Other Neocortical Pathologies |  |  |  |  |  |  |
| alpha-synuclein | - | - | - | - | - | - |
| TDP-43 | - | - | - | - | - | - |
| Used in: |  |  |  |  |  |  |
| Cryo-EM | + | + |  | + | + | + |
| IP | + | + | + | + | + | + |
| Mouse experiments | + | + |  |  | + | + |
| Fixed tissue staining |  | + |  | + | + |  |

**Extended Data Table 2.** Extracted brain tau lysates protein quantification.

| | Total Protein Concentration ($\mu\text{g}/\mu\text{L}$) | Tau Concentration ($\mu\text{g}/\mu\text{L}$) | Tau Purity (%) |
| --- | --- | --- | --- |
| **Mildly sonicated lysates (sark pel)** |  |  |  |
| AD Case 1 | 8.51 | 0.31 | 3.7 |
| AD Case 2 | 11.8 | 1.06 | 9.0 |
| VT Case 1 | 8.89 | 0.91 | 10.3 |
| VT Case 2 | 12.7 | 0.63 | 5.0 |
| VT Case 3 | 16.2 | 0.29 | 1.8 |
| **Extensively sonicated lysates (final sup)** |  |  |  |
| AD Case 1 | 4.82 | 0.57 | 11.8 |
| AD Case 2 | 3.29 | 0.74 | 22.5 |
| AD Case 3 | 2.88 | 0.47 | 16.3 |
| VT Case 1 | 3.33 | 0.80 | 24.1 |
| VT Case 2 | 12.8 | 1.07 | 8.4 |
| VT Case 3 | 10.5 | 1.34 | 12.8 |

**Extended Data Table 3.**
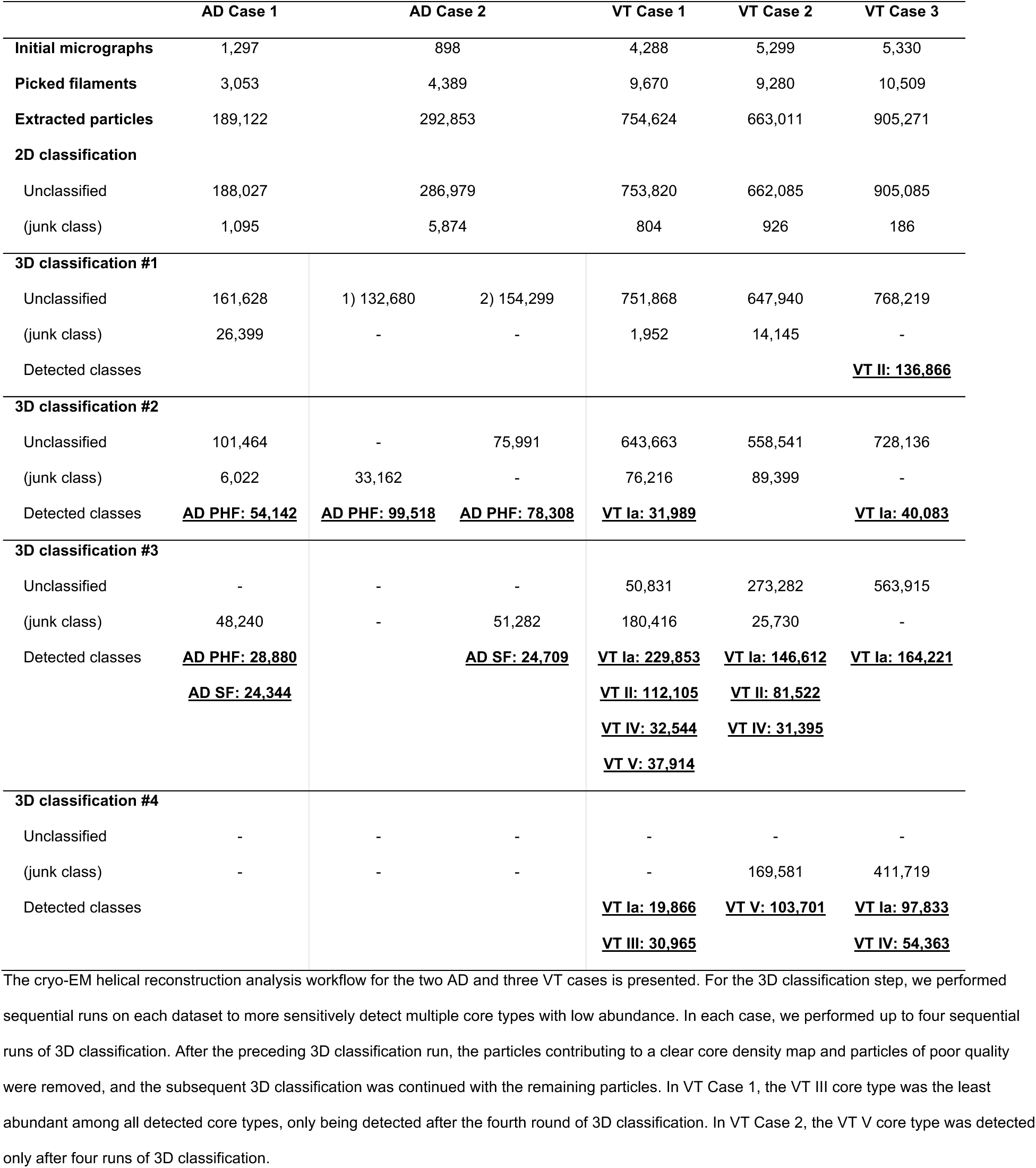
Cryo-EM workflow for AD and VT tau filament cores.

**Extended Data Table 4.**
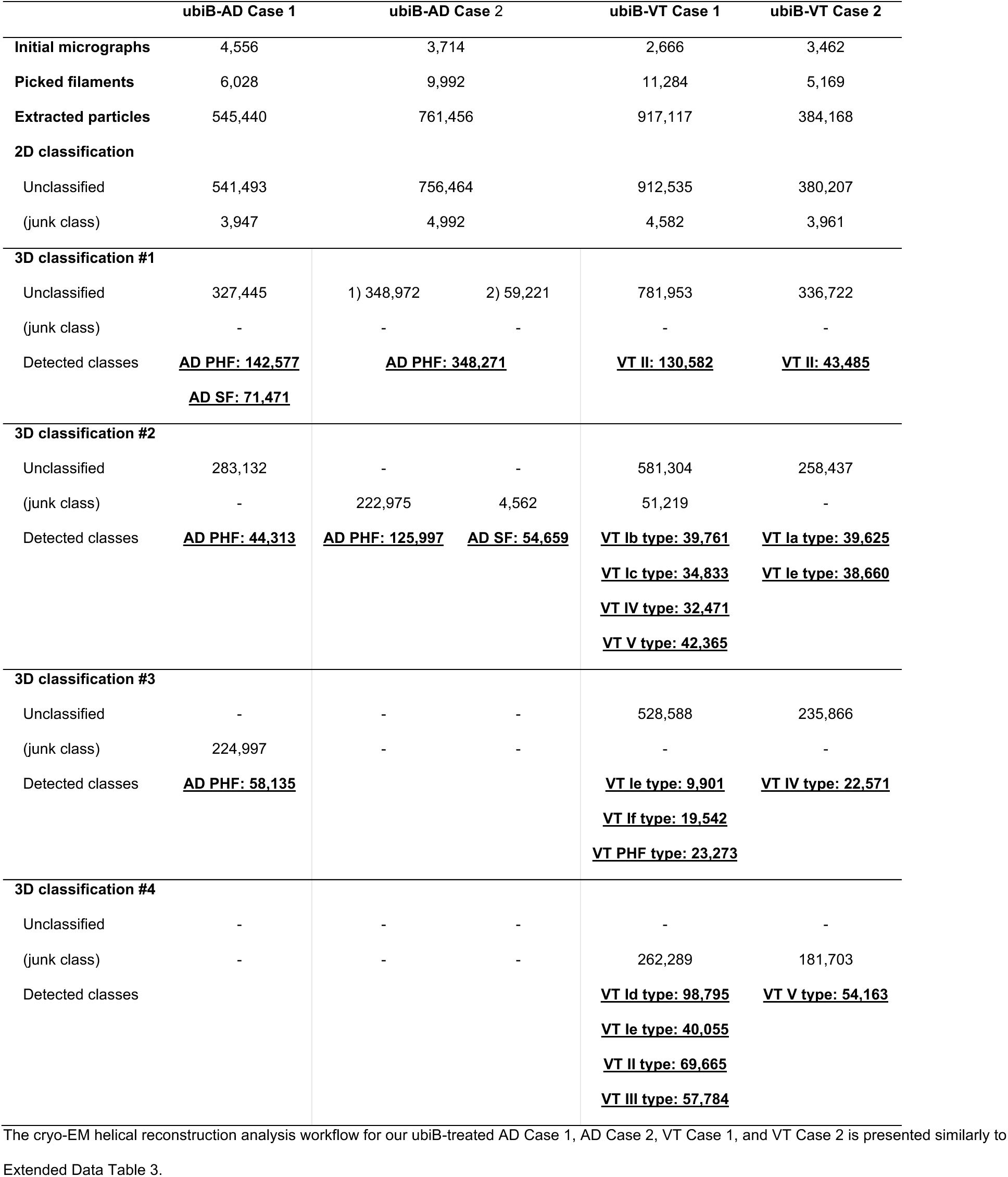
Cryo-EM workflow for ubistatin B-treated AD and VT tau filament cores.

|  | ubiB-AD Case 1 | ubiB-AD Case 2 |  | ubiB-VT Case 1 | ubiB-VT Case 2 |
| --- | --- | --- | --- | --- | --- |
| Initial micrographs | 4,556 | 3,714 |  | 2,666 | 3,462 |
| Picked filaments | 6,028 | 9,992 |  | 11,284 | 5,169 |
| Extracted particles | 545,440 | 761,456 |  | 917,117 | 384,168 |
| 2D classification |  |  |  |  |  |
| Unclassified | 541,493 | 756,464 |  | 912,535 | 380,207 |
| (junk class) | 3,947 | 4,992 |  | 4,582 | 3,961 |
| 3D classification #1 |  |  |  |  |  |
| Unclassified | 327,445 | 1) 348,972 | 2) 59,221 | 781,953 | 336,722 |
| (junk class) | - | - | - | - | - |
| Detected classes | <u>AD PHF: 142,577</u><br><u>AD SF: 71,471</u> | <u>AD PHF: 348,271</u> |  | <u>VT II: 130,582</u> | <u>VT II: 43,485</u> |
| 3D classification #2 |  |  |  |  |  |
| Unclassified | 283,132 | - | - | 581,304 | 258,437 |
| (junk class) | - | 222,975 | 4,562 | 51,219 | - |
| Detected classes | <u>AD PHF: 44,313</u> | <u>AD PHF: 125,997</u> | <u>AD SF: 54,659</u> | <u>VT Ib type: 39,761</u><br><u>VT Ic type: 34,833</u><br><u>VT IV type: 32,471</u><br><u>VT V type: 42,365</u> | <u>VT Ia type: 39,625</u><br><u>VT Ie type: 38,660</u> |
| 3D classification #3 |  |  |  |  |  |
| Unclassified | - | - | - | 528,588 | 235,866 |
| (junk class) | 224,997 | - | - | - | - |
| Detected classes | <u>AD PHF: 58,135</u> |  |  | <u>VT Ie type: 9,901</u><br><u>VT If type: 19,542</u><br><u>VT PHF type: 23,273</u> | <u>VT IV type: 22,571</u> |
| 3D classification #4 |  |  |  |  |  |
| Unclassified | - | - | - | - | - |
| (junk class) | - | - | - | 262,289 | 181,703 |
| Detected classes |  |  |  | <u>VT Id type: 98,795</u><br><u>VT Ie type: 40,055</u><br><u>VT II type: 69,665</u><br><u>VT III type: 57,784</u> | <u>VT V type: 54,163</u> |

**Extended Data Table 5.**
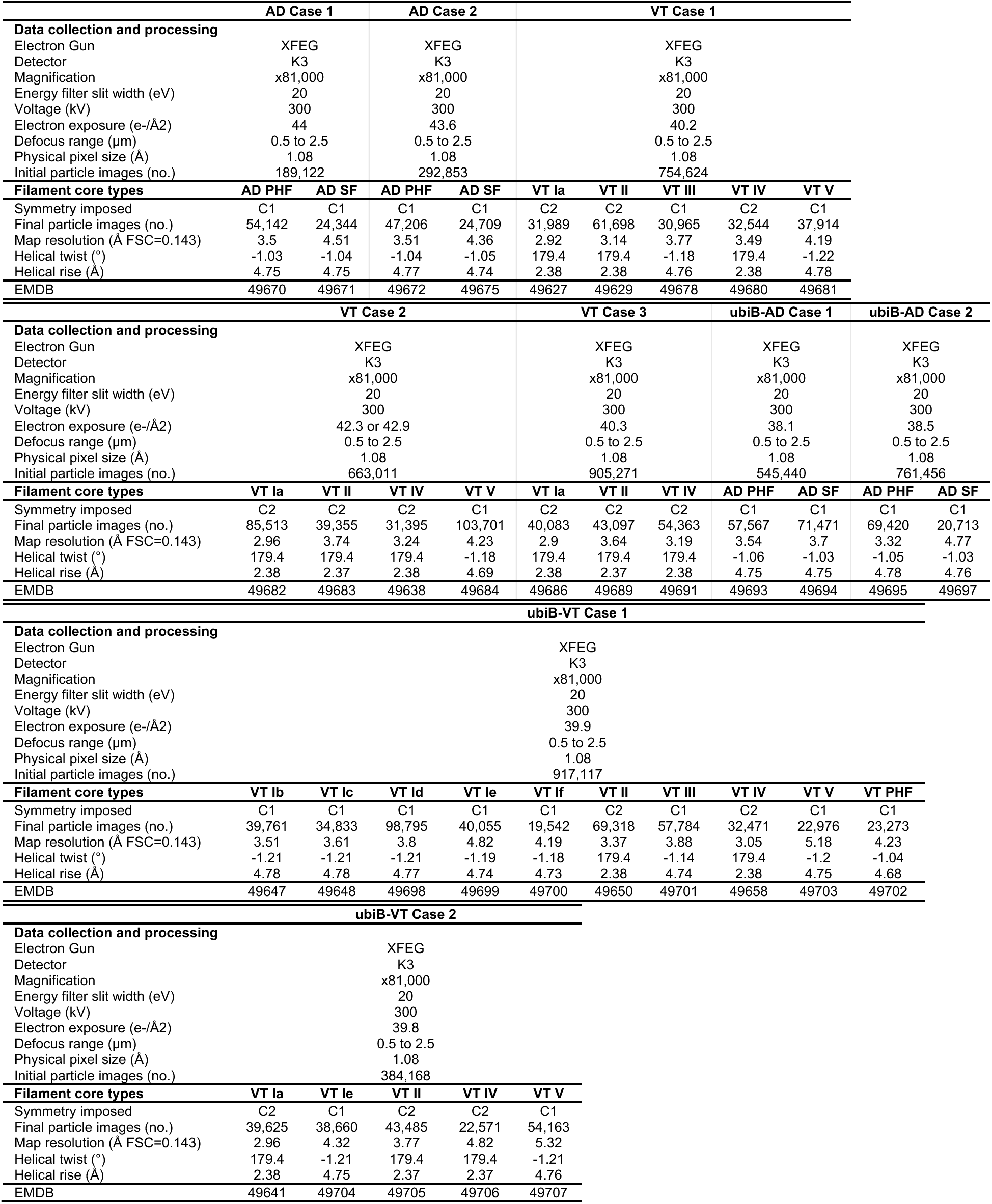
Cryo-EM data collection statistics.

**Extended Data Table 6.** Cryo-EM model refinement and validation statistics.

|  | VT Ia<br>VT Case 1 | ubiB-VT Ia<br>VT Case 2 | VT Ib<br>VT Case 1 | VT Ic<br>VT Case 1 | VT II<br>VT Case 1 | ubiB-VT II<br>VT Case 1 | VT IV<br>VT Case 2 | ubiB-VT IV<br>VT Case 1 |
| --- | --- | --- | --- | --- | --- | --- | --- | --- |
| **Model Refinement** |  |  |  |  |  |  |  |  |
| Model resolution (Å FSC=0.5) | 2.99 | 3.03 | 3.85 | 3.72 | 3.25 | 3.53 | 3.32 | 3.03 |
| Map sharpening B factor (Å<sup>2</sup>) | -31.34 | -38.74 | -56.80 | -70.89 | -65.15 | -63.67 | -55.17 | -36.26 |
| Model composition |  |  |  |  |  |  |  |  |
| Non-hydrogen atoms | 10206 | 10332 | 10332 | 10332 | 10206 | 10332 | 10206 | 10332 |
| Protein residues | 1350 | 1350 | 1350 | 1350 | 1350 | 1350 | 1350 | 1350 |
| Waters | 0 | 0 | 0 | 0 | 0 | 0 | 0 | 0 |
| B factors (Å<sup>2</sup>) |  |  |  |  |  |  |  |  |
| Protein | 58.92 | 48.19 | 91.12 | 70.46 | 54.23 | 28.09 | 59.96 | 48.19 |
| Waters | - | - | - | - | - | - | - | - |
| R.m.s. deviations |  |  |  |  |  |  |  |  |
| Bond lengths (Å) | 0.005 | 0.007 | 0.004 | 0.004 | 0.006 | 0.005 | 0.004 | 0.007 |
| Bond angles (°) | 0.956 | 1.064 | 0.847 | 0.792 | 0.857 | 0.933 | 0.657 | 1.064 |
| Validation |  |  |  |  |  |  |  |  |
| MolProbity score | 2.15 | 2.74 | 2.00 | 2.06 | 2.01 | 2.83 | 1.49 | 2.74 |
| Clashscore | 13.31 | 21.92 | 13.19 | 7.69 | 11.44 | 17.60 | 6.37 | 21.92 |
| Ramachandran plot |  |  |  |  |  |  |  |  |
| Favored (%) | 94.52 | 90.41 | 94.52 | 97.26 | 93.38 | 91.78 | 97.26 | 90.41 |
| Allowed (%) | 5.48 | 9.59 | 5.48 | 2.74 | 6.62 | 6.85 | 2.74 | 9.59 |
| Outliers (%) | 0 | 0 | 0 | 0 | 0 | 1.37 | 0 | 0 |
| PDB | 9NPK | 9NQ4 | 9NQB | 9NQC | 9NPQ | 9NQE | 9NQ0 | 9NQI |

**Extended Data Table 7.**
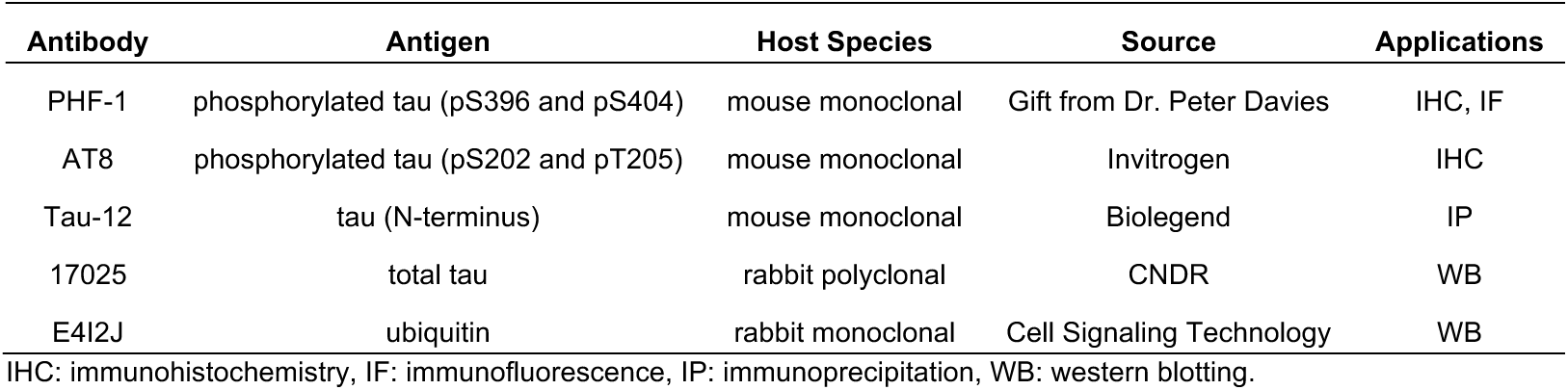
Primary antibodies used in this study.

## Supplementary information

**Supplementary Video 1. Shifting of VT I group filament cores**

The atomic models for VT I group filaments are presented sequentially in two different ways. First, atomic models with the structured tau filament in white are presented, with additional red or blue coloring based on electrostatic potential (corresponding to negative and positive, respectively). These are overlaid with their cognate density maps corresponding to untreated VT Ia, ubiB-treated VT Ia (VT Case 2), and VT Ib through VT If (VT Case 1) where the additional densities adjacent to the structured core are colored yellow. Second, successive line models of the structured tau core backbone are shown followed by their overlay.

